# Uptake and survival of *Leishmania amazonensis* in *Acanthamoeba castellanii*: An infection organism model?

**DOI:** 10.1101/2024.12.22.627952

**Authors:** Leonardo Fernandes Geres, Pedro Henrique Gallo Francisco, Diullia de Andrade Machado, Francisco Breno S. Teófilo, Marcelo Brocchi, Selma Giorgio

## Abstract

Leishmaniasis is a neglected tropical disease. Parasite strategies and the evaluation of drugs and vaccines are inferred from studies carried out using mouse models and macrophages. The development of model organisms with no ethical restrictions will contribute to our knowledge of leishmaniasis. *Acanthamoeba castellanii*, a free-living protozoan, is known to interact with various microorganisms. In this study, the interaction between the amoeba *A. castellanii* and the trypanosomatid *Leishmania amazonensis* was investigated by combining quantitative kinetics analysis, optical, fluorescence, electronic, confocal, and live video microcopy. We sought to standardize protocols for the co-culture; the optimal experimental conditions were: RPMI medium + 10% SFB at 26°C*. L. amazonensis* invades *A. castellanii* through its acanthopods, and the promastigotes interact with the trophozoites via their flagellum, which also occurs when parasites infect mammalian macrophages. The forms of *L. amazonensis* inside the amoeba become rounded and lose their flagellum; they are similar to amastigotes. These round forms were isolated from trophozoites after 3 h of co-culture and differentiated into promastigotes, demonstrating their viability inside amoeba. The percentage of amoebas with *L. amazonensis* was reduced overtime. Thus, considering that *A. castellanii* can clear *Leishmania*, this interaction could serve as an effective model of cellular leishmanicidal mechanisms.

## 1. INTRODUCTION

Leishmaniasis is designated by the World Health Organization as a neglected tropical disease (WHO, 2024). This parasitic disease is a global public health concern associated with poor communities (Burza et al., 2018). It is estimated that approximately 350 million people are at risk of infection by this parasite, and an average of 1 million new cases are reported each year (WHO, 2024). Epidemiological data have shown that the incidence of this disease has considerably increased in urban areas (Takele et al., 2022).

Leishmaniasis is transmitted through the blood meal of infected female sandflies of the genera *Phlebotomus* (Africa, Europe, and Asia) and *Lutzomyia* (mainly in South and Central America) (Serafim et al., 2021). After replication, the promastigotes in the sandfly digestive tract differentiate into the infective form, the metacyclic promastigotes. These are regurgitated during the blood meal, together with the insect saliva, into the dermis of the mammalian host (Serafim et al., 2021). The inoculated saliva attracts phagocytic cells to the bite site, and the metacyclic promastigotes, when phagocytosed by the host cells of the mononuclear phagocyte system, transform into amastigotes and begin to proliferate (Serafim et al., 2021).

Among the different clinical manifestations of the disease, cutaneous leishmaniasis is the most common; this form is painless and is usually located in exposed areas of the skin, with an erythematous base and well-defined edges (Reithinger et al., 2007; Alvar et al., 2012). Mucocutaneous leishmaniasis destroys mucosal tissue, mainly in the nasopharyngeal region (Reithinger et al., 2007). Visceral leishmaniasis, which causes swelling of the abdomen, due to the enlargement of the liver and spleen, is the most serious form of leishmaniasis and can be lethal if left untreated (Alvar et al., 2012). The treatments available for leishmaniasis include meglumine antimoniate, amphotericin B, pentamidine, and miltefosine(Croft and Olliaro, 2011; Uliana et al., 2018). Except for miltefosine, all of these drugs are administered parenterally, which makes treatment difficult to adhere to. Moreover, there is a limited number of drugs available, and these are associated with, problems such as severe side effects and high cost of treatment (Uliana et al., 2018). Furthermore, no vaccines are currently available to prevent leishmaniasis in humans (Alvar et al., 2012; Burza et al., 2018). Owing to the lack of adequate attention, drug screening is still necessary. In order to develop more effective treatments, it is essential to develop adequate models that can mimic the cellular environment and the life cycle of the parasite.

Organism models must necessarily be representative of the desired system and be easier to study than the modeled target (i.e., humans), in each field of research, there are specific criteria in selecting model organisms(Leonelli and Ankeny, 2013; Montagnes et al., 2012). The complex relationships between *Leishmania* and its escape mechanisms, infection, metabolic pathways, drug candidates, and cytotoxicity have been well studied in cellular systems (Brioschi et al., 2022). The parasite strategies, microbicidal functions, immunomodulation, and the evaluation of effectiveness of drugs and vaccines have been inferred from studies carried out with primary murine macrophages (peritoneal and bone marrow-derived) and immortalized cell lines (Hendrickx et al., 2012; Terreros et al., 2019). However, all these models have limitations, and there is no single model that mimics all aspects of infection and pathogenesis observed in humans. It is expected that we will have different models that provide different answers, considering that in humans, the development of the parasite is very uncertain and dependent on several factors such as immunological factors, host microbiota, and *Leishmania* species. Therefore, the development of more models can help in understanding the diversity of the pathogen-host relationship of this parasite and contribute to this diverse web of possibilities.

There is a wide diversity of protozoa that are characterized by short generation time and easy storage, handling, and identification (morphological, genetic, and biochemical data), as well as phenotypic stability. These organisms are suitable for interdisciplinary studies, have no ethical restrictions, are more economical to maintain than are vertebrates all traits that make them suitable as model organisms, thus facilitating physiological studies and increasing interest in research on protozoa(Montagnes et al., 2012).

*Acanthamoeba castellanii*, is a ubiquitous and free-living protozoan that reproduces through mitosis. Trophozoites are often associated with biofilms(Geres et al., 2024). Cysts can be transmitted through the air (winds and dust storms), such as long-distance ventilation ducts (Lacerda and Lira, 2021).

*A. castellanii* is known to interact with various microorganisms, including bacteria (e.g. *Legionella pneumophila*), fungi (e.g. *Trichophyton rubrum*), viruses (e.g. Yaravirus), and protozoa (e.g. *Toxoplasma gondii* and *Cryotosporidium parvum*) (de Faria et al., 2020; Fukaya et al., 2023; Mungroo et al., 2021; Winiecka-Krusnell et al., 2009). In this interaction, the amoebas may act as a predator, a transmission vehicle when the microorganism lives within the amoebas but does not multiply, or a reservoir when the microorganism proliferates inside the amoebas (Mungroo et al., 2021). Interestingly, these characteristics are shared with macrophages. In addition, *A. castellanii* and mammalian macrophages share several similarities in structure, cellular physiology, presence of digestive vacuoles, phagotrophic capacity, and ability to eliminate microorganisms. For example, oxidative attack in amoebas is similar to what is observed in macrophages in the presence of reactive oxygen species (ROS) and nitric oxide (NO) (Mungroo et al., 2021; Rayamajhee et al., 2021). The hypothesis that *A. castellanii*, in some evolutionary period, may have been a host of *Leishmania* and “trained” it to evade phagocytes and consequently the innate immune system, was suggested, considering that the two protists have been present on Earth for millions of years(Ahmed, 2014).

There are only a few reports on the interaction between *Leishmania* and amoebas and, the details of their relationship have not been fully explored (Campo-Aasen et al., 1988; Santos et al., 2024; Winiecka-Krusnell et al., 2009). Campo-Aasen et al. suggested based on analysis of electron microscopy images that the interaction of *A. castellanii* with *Leishmania* leads to the destruction of the amoeba(Campo-Aasen et al., 1988), and Santos et al., analyzing optical microscopy images, concluded that *Leishmania* subverts amoeba functions (Santos et al., 2024). In the present study, we combined quantitative kinetics analyses and optical, fluorescence, electronic, confocal, and live video microscopy to better understand *in vitro* interactions between *A. castellanii* and *L. amazonensis*.

## 2. MATERIAL AND METHODS

### 2.1 Parasite culture

*L. amazonensis* (MHOM/BR/73/M2269) promastigotes were cultivated in RPMI-1640 medium (Sinergia, Campinas, Brazil) supplemented with 10% fetal bovine serum (FBS) (Vitrocell, Freiburg, Germany) and 50µg/m gentamicin (Sigma-Aldrich) at pH 7.4 in 25-cm^2^ cell culture flasks at 26°C (Bacteriológica SL 101– SOLAB)(de Mesquita Barbosa et al., 2015). The promastigote forms of *L. amazonensis* expressing green fluorescent protein (*L. amazonensis*-GFP) (strain MHOM/BR/75/Josefa) were maintained in RPMI culture medium + 10% FBS and periodically selected with 250µg/mL geneticin (G418) (Sigma-Aldrich, St. Louis, MO, USA), grown in 25-cm^2^ cell culture flasks at 26°C (Costa et al., 2011).

*A. castellanii* trophozoites (ATCC 30010) were kindly provided by Prof. Dr. Cristina Elisa Alvarez Martinez from the Department of Genetics, Evolution, Microbiology, and Immunology at the Institute of Biology, Universidade Estadual de Campinas (Unicamp). Trophozoites were cultivated in culture flasks in axenic peptone-yeast extract medium + 0.1 M glucose (PYG-W), containing 300 µg/mL streptomycin and 100 µg/mL ampicillin in 25-cm^2^ cell culture flasks at 26°C (Winiecka-Krusnell et al., 2009).

### 2.2 Parasite proliferation curves

Proliferation curves of promastigotes cultured in RPMI medium + 10% FBS, PYG-W medium or PYG-W medium + 10% FBS at 26°C were obtained. Promastigotes were cultured in RPMI medium + 10% SFB, in PYG-W medium or in PYG-W medium + 10% SFB at 26°C, 34°C or at 37 °C. In both cases, 5 × 10^5^ parasites/mL were cultured in 6-well plates at a final volume of 5 mL in the respective medium. At 24-h intervals, parasites were quantified using a Neubauer chamber for at least 7 days, in triplicate wells(de Faria et al., 2020; Nunes et al., 2013).

### 2.3 Uptake assays

Assays were performed in 24-well plates containing 13-mm diameter coverslips. The interaction of *A. castellanii* trophozoites with *L. amazonensis* promastigotes or polystyrene microbeads(diameter 2 μm) fluorescein isothiocyanate (FITC) immobilized on microbeads (Thermo Fisher, MA, USA) was carried out in PYG-W medium + 10% FBS or in RPMI medium + 10% FBS. The plates were incubated at 26°C or 34°C. The parasites were added to the wells at ratios of 1:10 and 1:20 (trophozoites: promastigotes). Microbeads were added to the wells at trophozoite-to-microbead ratio of 1:10. The coverslips were removed from the wells at 3, 24, 48, and h ofco-cultivation, washed in PBS, fixed in absolute methanol, and stained with Giemsa (Merck, Darmstadt, Germany) or Rapid Panoptic Stain (Laborclin, Pinhais, Brazil)(de Faria et al., 2020; Winiecka-Krusnell et al., 2009). The slides were analyzed using a common optical microscope (Primo Star Zeiss), and images were captured using AxioVision 4.8. Image adjustment (brightness, contrast, and scale) and analysis was performed on ImageJ (Garajová et al., 2019; Karaś et al., 2018). Three independent experiments were performed in triplicate. The intact and adherent *A. castellanii* trophozoites were counted in 20 representative random fields to measure the viability of *A. castellanii* trophozoites. In parallel, counts of 200 trophozoites were made, and the following were quantified: percentage of amoebaswith *L. amazonensis*, average number of intracellular forms per trophozoite, percentage of amoebas containing microbeads, and average number of microbeads per trophozoite. All counts were made at 1,000× magnification using optical and fluorescence microscope (de Faria et al., 2020; de Mesquita Barbosa et al., 2015).

### 2.4 Viability of L. amazonensis internalized by A. castellanii trophozoites

Following the co-cultivation assays described previously, intracellular forms were recovered by lysing the trophozoites after 3 h of interaction between *A. castellanii* trophozoites and *L. amazonensis* promastigotes in RPMI medium + 10% FBS or PYG-W at ratio of 1:10 or 1:20 (trophozoite: promastigotes). The cultures were washed three times with PBS 1× and lysed with 0.04% sodium dodecyl sulfate (SDS) in PBS through seven passages of the contents using a 30-G needle in sterile 1-mL syringes (de Faria et al., 2020). The suspension was centrifuged (100 *g*) for 10 min and the supernatant was collected and centrifuged again (800 *g*) for 10 min. The pellet was resuspended in RPMI medium + 10% FBS(de Faria et al., 2020). The recovered intracellular forms were seeded in 24-well plates with RPMI medium + 10% FBS for differentiation into promastigotes at 26°C and counted in a Neubauer chamber for a period of 10 days (de Faria et al., 2020; Mendes et al., 2022).

### 2.5 Fluorescence and confocal microscopy

*A. castellanii* with *L. amazonensis*-GFP or microbeads were co-cultured in 24-well plates with 13-mm diameter coverslips in either PYG-W medium supplemented with 10% SFB, or in RPMI medium supplemented with 10% FBS. They were plated in a 1:10 ratio (trophozoites: promastigotes or microbeads) and incubated at 26°C. Parasites were fixed in 4% paraformaldehyde after the coverslips were removed, then washed with PBS and permeabilized with triton-X (0.3%) for 1 min. The coverslips were then stained with Celltracker (diluted 1:1,000 in PBS 1X) (cytoplasm) (Thermo Fisher) for 30 min, followed by 4’,6-diamidino-2-phenylindole(DAPI diluted 1:1000 in PBS 1x) (nucleus cell) (Thermo Fisher) for 40 min, and washed with PBS. Lastly, the coverslips were placed on 4 µL of Vecta Shield mounting medium (Garajová et al., 2019; Karaś et al., 2018). The slides were then analyzed under a fluorescence microscope and an confocal microscope with Airyscan mode (Zeiss LSM880 Airyscan AG, Germany), was used for DAPI excitation (emission filter 450/40 nm), for FITC excitation (emission filter 510/20 nm), and for CellTracker Red excitation (emission filter 620/30 nm), at 63× oil immersion objective, with zoom 1x(Albuquerque et al., 2019). Images were captured using ZEN (Carl Zeiss Microscopy GmbH) software. Image adjustment (brightness, contrast, and scale) was performed using the Fiji 3D Script/Vaa3D software.

### 2.6 Live microscopy

The analysis of the real-time interaction of *A. castellanii* trophozoites with *L. amazonensis*-GFP promastigotes was carried out in a 16-well chamber slide system (LabTek, São Paulo, BR), in RPMI medium supplemented with 10% FBS, in a ratio of 1:10 (trophozoites: promastigotes) and incubated at 26°C for 24 h. The recordings were performed under an inverted microscope confocal with Airyscan mode (Zeiss LSM880 Airyscan AG, Germany), at 20×oil immersion objective(Hochstrasser and Hilbi, 2019; Mengue et al., 2017). Images were captured using ZEN (Carl Zeiss Microscopy GmbH) software. Image adjustment (brightness, contrast, and scale) was performed using the Fiji software.

### 2.7 Scanning Electron Microscopy

Topological changes due to parasite interactions were examined using scanning electron microscopy. To this end, 1 × 10^5^ *A. castellani* trophozoites and 1 × 10^6^ *L. amazonensis* promastigotes (proportion of 1:10, trophozoite: promastigotes) were plated on 13-mm coverslips and, after 2 h, fixed in 2.5% glutaraldehyde (Electron Microscopy Sciences, Hatfield, PA) in 0.1M PBS. After washing with PBS, the samples were postfixed in 1% osmium tetroxide (1h) (Electron Microscopy Sciences, Hatfield, PA), sequentially dehydrated in ethanol, the samples were washed, then brought to the Critical point dryer (Balzers CPD-030), and covered with Au using Sputter Coater (Balzers SCD-050). Lastly, the samples were analyzed using a scanning electron microscope (JEOL JSM-5800LV), operating at a standard accelerating voltage of 10 kV(Albuquerque et al., 2019; Fakae et al., 2020; Kathuria et al., 2014). Image adjustment (brightness, contrast, and scale) was performed using the Fiji software.(Albuquerque et al., 2019; Fakae et al., 2020; Kathuria et al., 2014).

### 2.8 Polymerase chain reaction (PCR)

DNA was extracted from both parasites (*A. castellanii* and *L. amazonensis*) using QIAamp DNA Mini Kit (Qiagen, Hilden, DE) with 5 × 10^6^ promastigotes and 5 × 10^6^ trophozoites for DNA amplification, the primers G6PDH-F (5’- CGYCTYCCAGACGCYTACGA-3’) and G6PDH-R (5’- AGCGGYGTGAAGATGCGCCA-3’) were used to detect a 110-bp fragment of the nuclear gene that encodes the enzyme G6PD (glucose 6-phosphate dehydrogenase) of *Leishmania* (Coser et al., 2020). Each reaction mix contained 1 µL of DNA target, 1 μL of primers, 6.25 µL of Gotaq (Hot Start Green Master Mix, Promega®, Brazil), and 50 µL of deionized water. The PCR conditions consisted of denaturation at 95 °C for 3 min, followed by 40 cycles of denaturation at 95 °C for 15 s, annealing at 60 °C for 30 s, and extension at 72 °C for 90 s. The samples were analyzed by electrophoresis on 1% agarose gels stained with ethidium bromide(Coser et al., 2020).

### 2.9 Statistical analysis

The experiments were repeated at least three times independently. The mean and standard deviation (SD) were calculated for all assays, and analysis of variance (ANOVA) was performed between the experimental situations, using the GraphPadPrism version 8.1 program. Statistical significance was verified by the ANOVA test; P values ≤ 0.05 were considered statistically significant.

## 3. RESULTS

### 3.1 Uptake assays: interaction between A. castellanii trophozoites and L. amazonensis promastigotes at 26°C and 34 °C in RPMI and PYG medium

We standardized the proliferation curves in RPMI and PYG mediafor both *L. amazonensis* and *A. castellanii* (Figure S1). PYG is the most commonly used medium for cultivating *A. castellanii*,and RPMI is commonly used for *L. amazonensis* (Latifi and Salimi., 2020; Oliveira-Filho et al., 2024). Both the parasites proliferated as previously described.

We then, analyzed the interaction between *A. castellanii* trophozoites and *L. amazonensis* promastigotes at 26°C in RPMI and PYG media. After 3 h of interaction, in RPMI medium at 26°Cat different ratios of trophozoite: promastigotes(1:10 and 1:20) the average percentage of amoebas containing *L. amazonensis* was 20.1% ± 6.4 and 25.1% ± 5.3, respectively (Figure 1A). The number of intracellular forms of *L. amazonensis* varied from 1.6 to2.8/amoeba. After 24 h of interaction (1:10 and 1:20, trophozoite:promastigotes), the percentages with *L. amazonensis* in RPMI medium were 11.1% ± 1.1 and 23.6% ± 2.5, respectively (Figure 1A). The number of intracellular forms of *L. amazonensis* varied between 1.1 and 2/amoeba (Figure 1B). At the timepoints of 48 and 72 h,no surviving *L. amazonensis* were evident within the *A. castellanii*.

**Figure 1.**
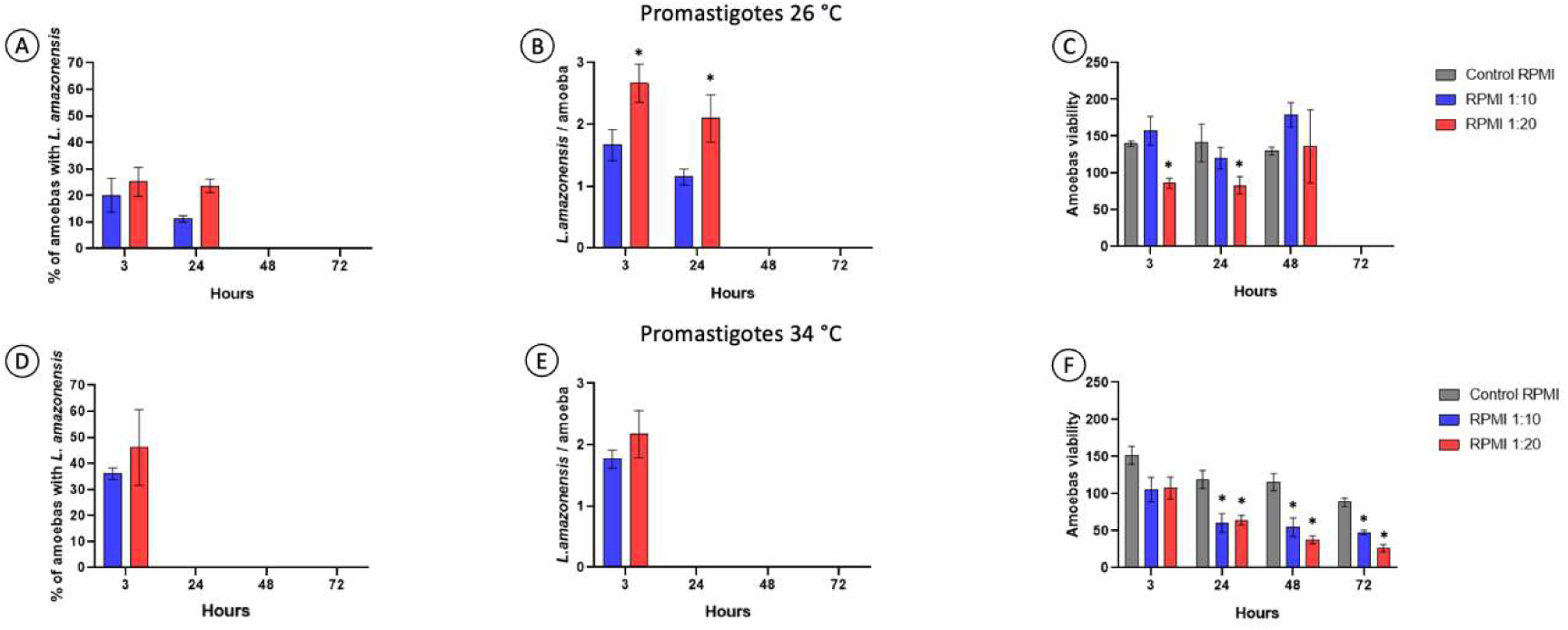
Interaction between *A. castellanii* trophozoites and *L. amazonensis* promastigotes. Trophozoites and promastigotes were maintained at 26°C (A, B and C) and 34°C (D, E and F) in RPMI + 10% SFB, 3, 24, 48 and 72 hours, with parasite ratios of 1:10 and 1:20 (trophozoite: promastigotes). Data are representative of three independent experiments, performed in triplicate and values are expressed in mean ± SD; *p ≤ 0.05 (ANOVA).

The viability of amoeba in RPMI medium (1:10, trophozoite:promastigotes) at 26°C remained statistically unchanged at 3, 24, and 48 h. However, in the case of the 1:20 treatment, there was an approximately 45% decrease in viability at the 3 and 24 h mark, compared with control group (only amoebas). No statistical changes were observed in this treatment group at 48 h (Figure 1C).

Through optical microscopy we confirmed that *L. amazonensis* promastigotes were able to interact with *A. castellanii* trophozoites *in vitro* through the flagellum, as occurs when promastigotes infect cells of the mononuclear phagocyte system; most of the “phagocytosed” forms of *L. amazonensis* became rounded and lost the flagellum (similar towhat occurs with amastigotes) (Figure 2, red arrow) at 3 and 24 h. Photomicrographs of the interaction kinetics indicated a decrease in the forms of *L. amazonensis* over time (Figure S2).

**Figure 2.**
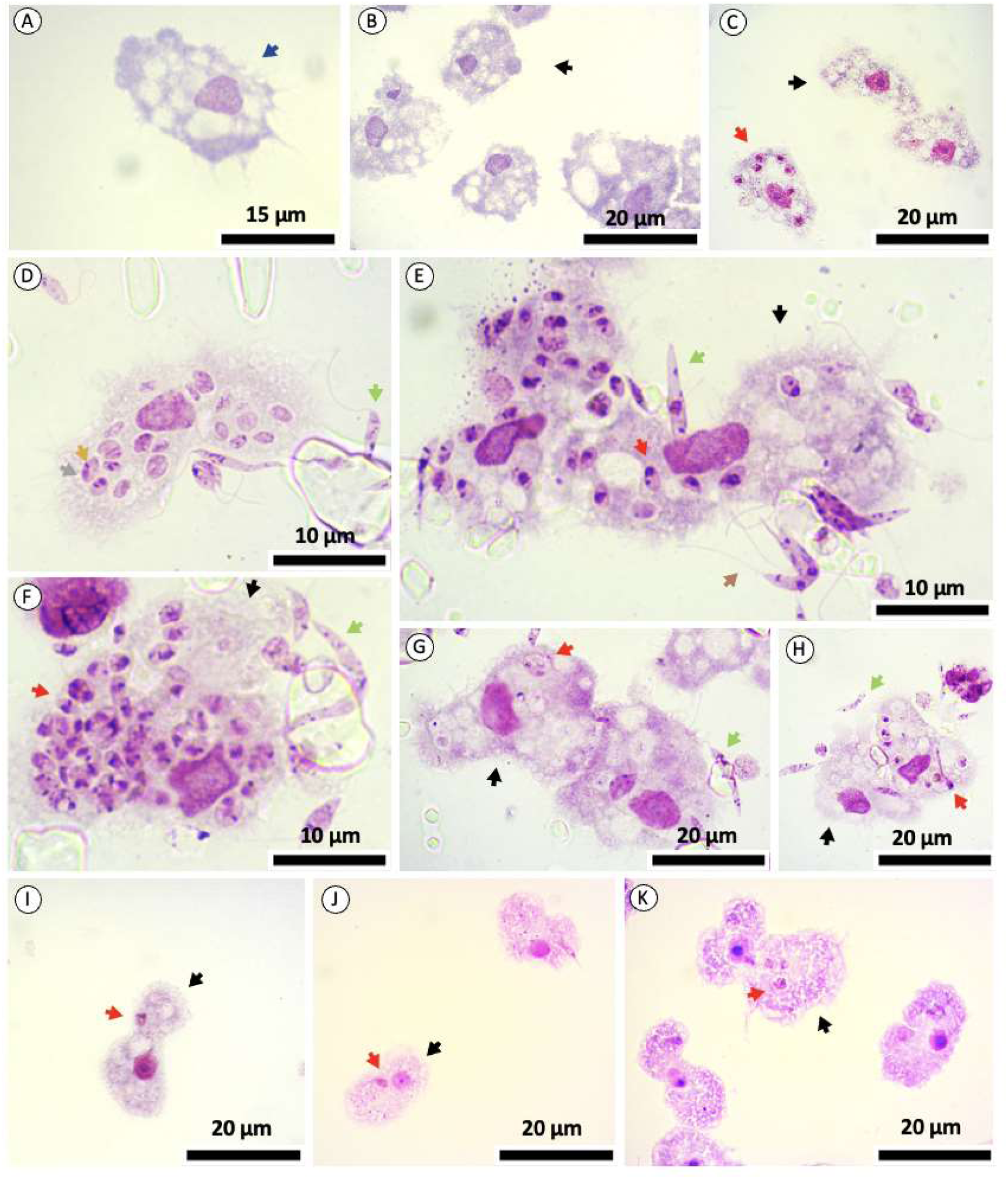
*A. castellanii* trophozoites in co-culture with *L. amazonensis* maintained for 3 hours. Red arrows: intracellular cells of *L. amazonensis*, black arrows: trophozoites, green arrows: promastigotes, blue arrows: trophozoite nucleus, gray arrows: *L. amazonensis* nucleus, yellow arrows: *L. amazonensis* kinetoplast. Cells were stained with Giemsa. Trophozoites (A-B); trophozoites in co-culture with *L. amazonensis* promastigotes maintained for 3 hours at 26°C (C-I); trophozoites in co-culture with *L. amazonensis* promastigotes maintained for 3 hours at 34°C (J-K).

We also evaluated the interaction between *A. castellanii* and *L. amazonensis* promastigotes in medium PYG at 26°C (Figure S3); however, we found that the RPMI medium at 26°C is preferable for this model of interaction, given that *L. amazonensis* promastigotes are more internalized in RPMI than in PYG medium (Figure 1A, Figure S3A). Because the percentage of amoebae within *L. amazonensis* is higher in RPMI medium (20.1% ± 6.4) than in PYG medium (5.3% ± 1.5) (Figure S3A), there were approximately 25% fewer amoebas with *L. amazonensis* than was the case in the RPMI medium (1:10, trophozoite:promastigotes). The RPMI medium appears to be the best environment for *Leishmania* culture.

With the aim of mimicking infection in macrophages, we analyzed the interaction between *A. castellanii* and *L. amazonensis* promastigotes at 34°C, as this is approximately the temperature of the host skin. After 3 h in RPMI medium, the average percentage of amoebas with *L. amazonensis* after treatment with 1:10 and 1:20 (trophozoites: promastigotes) was 36% ± 2.29 and 46.1% ± 14.4, respectively (Figure 1D). These values were close to those found in co-cultures at 26°C (20%, 30%) (Figure 1A). The number of intracellular forms of *L. amazonensis* ranged from 1.4 to 3 (Figure 1E) and were greater than in co-cultures at 26°C (1-2) (Figure 1B). At the timepoints of 24, 48 and 72 h, the *L. amazonensis* was unable to survive within the *A. castellanii* (Figure 1E). It should be noted that high infection rates were observed with promastigote and mammalian macrophage cell lines cultures (data not shown).

The viability of amoebas in RPMI medium at 34°C was reduced by approximately 50% after 24 h (1:10 and 1:20, trophozoite:promastigotes), when compared to control (Figure 1F). The viability of amoeba was approximately 20% lower than in the RPMI medium at 26°C (Figure 1C).

We also evaluated the interaction between *A. castellanii* and *L. amazonensis* promastigotes in PYG medium at 34°C (Figure S2). However, the percentage of amoebae with *L. amazonensis* in both RPMI and PYG media was 35% after 3 h (Supplementary 3A). At 24, 48, and 72 h, *L. amazonensis* was unable to survive within *A. castellanii* (Figure S2).

*L. amazonensis* amastigotes isolated from murine lesions were tested in co-cultures, and the results indicated that the interaction and internalization of this parasite form also occurred in *A. castellanii* trophozoites (data not shown). We also tested the interaction between *L. amazonensis* promastigotes and cysts, and the results indicated that the promastigotes were not internalized by cysts (data not shown).

In conclusion, we confirmed that to obtain maximum rate of internalization of *L. amazonensis* in the amoeba hosts, the RPMI medium at 26°C is preferable to PYG medium.

### 3.2 Viability of internalized forms of L. amazonensis

Intracellular forms of *L. amazonensis* were successfully isolated from *A. castellanii* trophozoites 3 h after co-culture. Their transformation into promastigotes and the rate of proliferation were followed for 10 days. Promastigotes were quantifiable after eight days in culture, confirming that the recovered parasites were viable (Figure S4). However, after 24 h of co-culture, no *L. amazonensis* were recovered from within the trophozoites after 24 h.

### 3.3 Uptake assay: microbeads

To evaluate the uptake capacity of the trophozoites, we used fluorescent microbeads. We observed that the amoebas were able to uptake the microbeads at 26°C or 34°C in either RPMI or PYG media (Figure S4). After 3 h of incubation, 40% to 50% of amoebas contained microbeads at both temperatures and media (Figure S5A). This ratio increased over time, reaching approximately 90% (Figure S5A). The average number of microbeads per amoeba varied between 2 and 5 at almost every timepoint (Figure S5B).

Amoeba viability decreased by approximately 25% over time at 26°C (Figure S4C). At 34°C the viability of the amoebae was approximately 20% lower when compared to the experimental groups at 26°C (Figure S5C).

### 3.4 Optical, fluorescence, confocal microscopy and live microscopy

We also confirmed that *L. amazonensis* is internalized by *A. castellani* by using *L. amazonensis*-GFP co-cultured with trophozoites (1:10, trophozoites:promastigotes). In this co-culture, approximately27% of amoebas were found to contain *L. amazonensis*, with an infection rate of 0.8-2.8/amoeba after 3 h, indicating that the interaction results are similar to what was obtained with the *L. amazonensis* wild-type strain (Figure 1). Using fluorescence microscopy, it was possible to observe acanthopods in the trophozoites as well as the interaction of promastigote forms with trophozoites through their flagellum (Figure 3). Through orthogonal visualization of the XYZ axes using a confocal microscope (Figure 4), we confirmed the observation that *L. amazonensis*-GFP was inside the amoeba and did not overlap it. This was corroborated by the 3D processing of 31 interaction planes (Figure S6 and S7, and Supple Movie S1). In addition, using time-lapse, we observed that the number of *L. amazonensis*-GFP was reduced after 6 and 24 h of interaction compared with 3 h, and confirmed that the amoeba phagocytosed *L. amazonensis*-GFP through its acanthopods (Movie S2, Movie S3).

**Figure 3.**
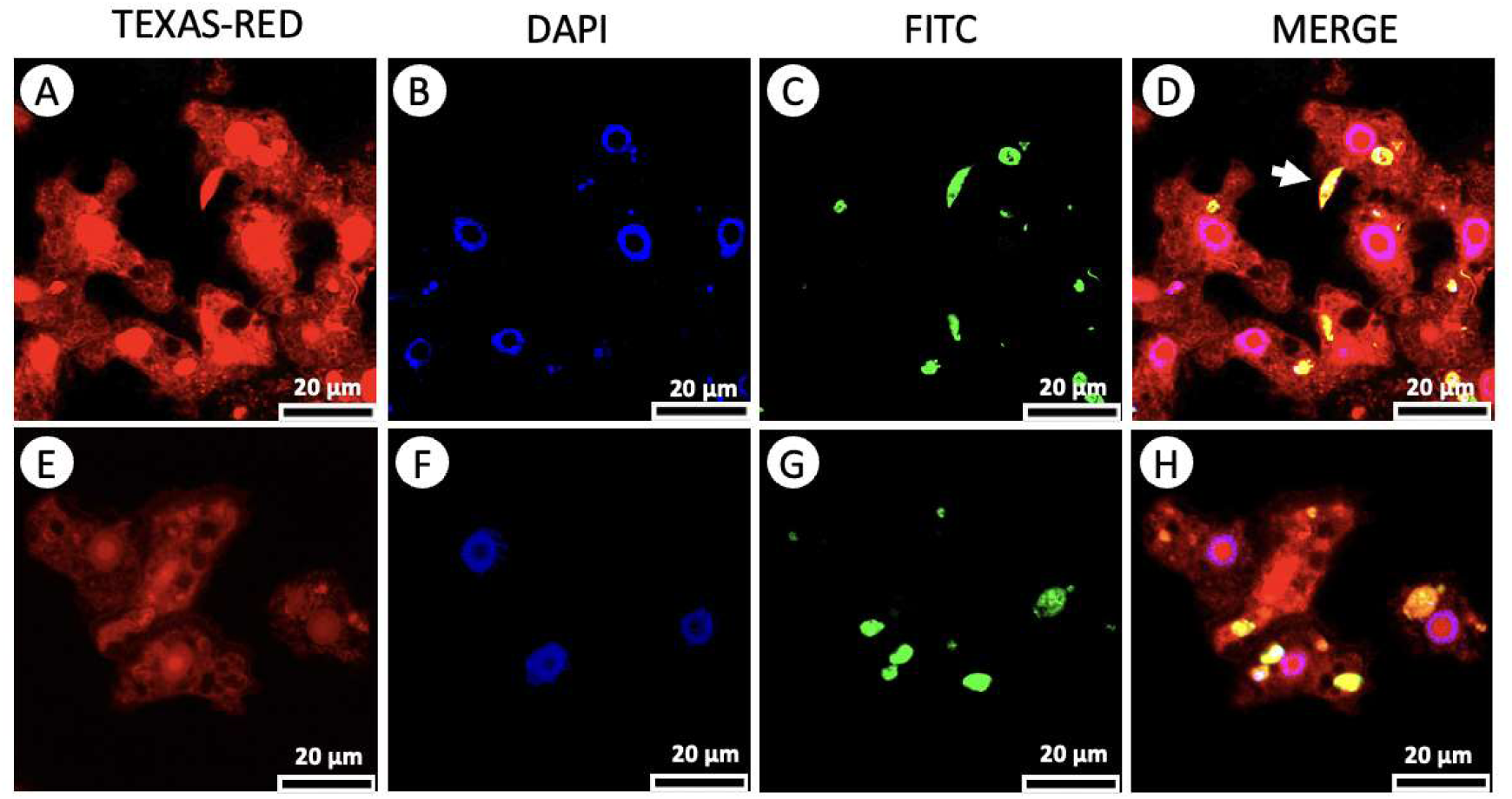
Confocal microscopy of parasite co-cultures stained with DAPI (nucleus) and Celttracker (cytoplasm). Celltracker (A and E). DAPI (B and F). FITC (*L. amazonensis*) (C and G). Merge (D and H). White arrow indicates promastigote interacting by flagellum with trophozoites.

**Figure 4.**
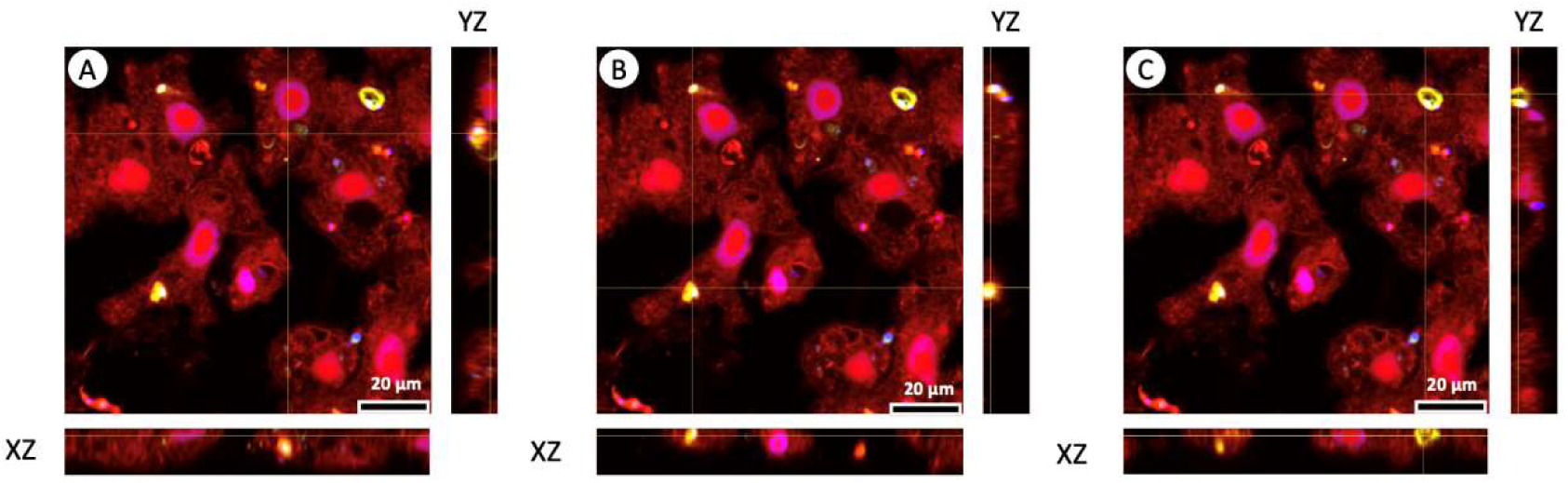
Orthogonal cutting confocal microscopy parasite co-cultures.

### 3.5 Topological analysis and parasite morphological changes in co-cultures

To topologically analyze the interaction of parasites, we performed scanning electron microscopy of *A. castellanii* trophozoites (control) and *L. amazonensis* (control) and compared them with trophozoites that interacted with promastigotes for 2 h (Figure 5A-D). We observed that the promastigote forms interacted with *A. castellanii* trophozoites, mostly through their flagellum. Interestingly, the surface of the trophozoites appeared to be more damaged during the interaction compared with the control trophozoites (Figure 5F, blue arrow).

**Figure 5.**
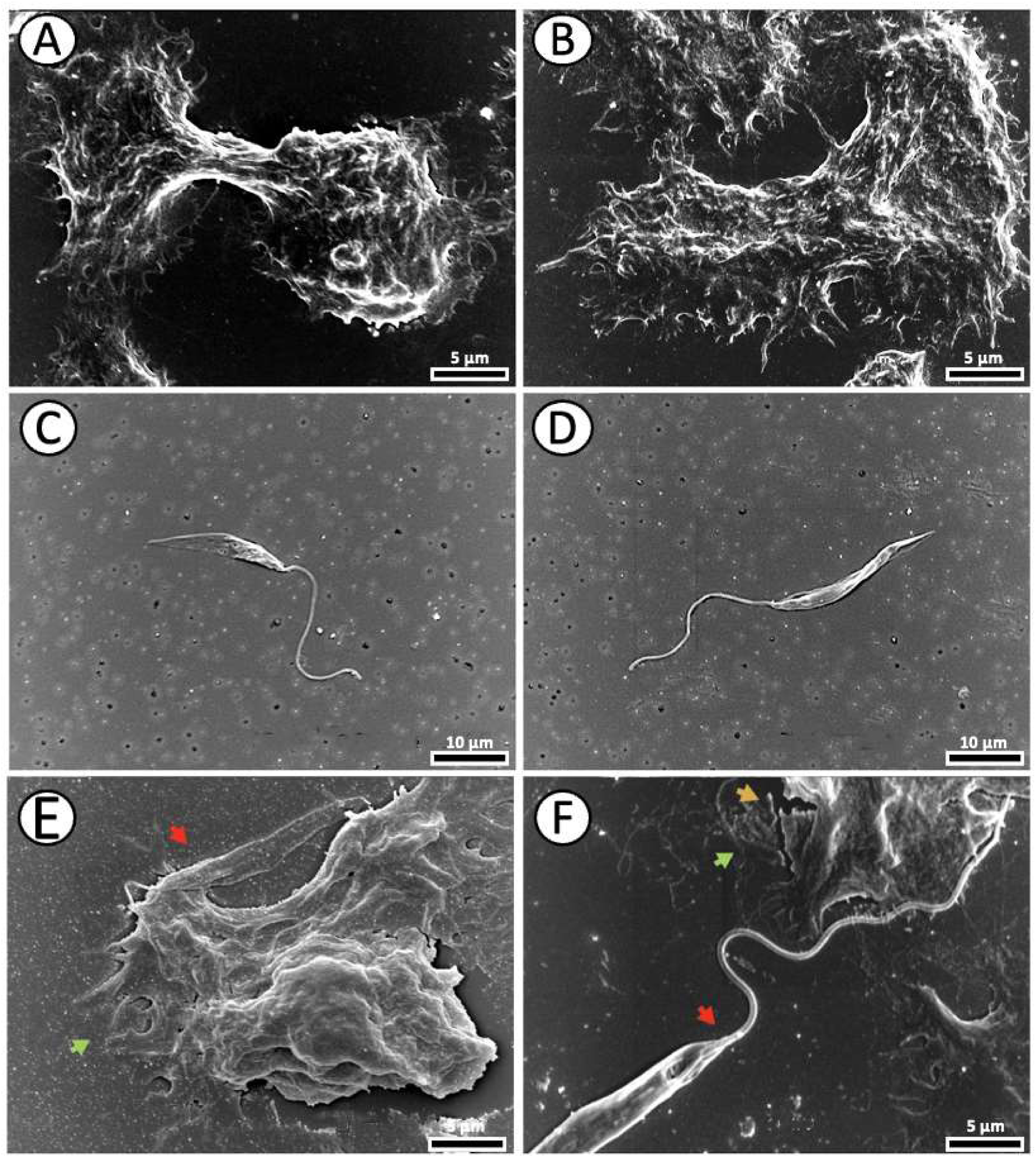
Scanning electron microscopy of *A. castellanii* trophozoites interacting with *L. amazonensis* promastigotes. (A, B) Trophozoites. (C, D) Promastigotes. (E, F) *A. castellanii* trophozoites interacting with *L. amazonensis* promastigotes. Red arrows indicate *L. amazonensis* promastigotes, green arrows indicate *A. castellanii* trophozoites, yellow arrow indicate surface damaged.

## 4. DISCUSSION

We have provided an understanding of the interaction between the amoeba *A. castellanii* and Trypanosomatidae *L. amazonensis* by combining quantitative kinetics analyses and optical, fluorescence, electronic, confocal, and live video microscopy. Our group also sought to standardize protocols for promastigote and trophozoite co-cultures at different temperatures, culture media, interaction ratios, and cultivation times with the best experimental conditions for analyzing the interaction between amoeba and *Leishmania* being in RPMI medium + 10% SFB at 26°C, co-culture ratio of 1:10 (trophozoites: promastigotes) and exposure time of 3-24 h after co-cultivation. In this experimental situation, *L. amazonensis* was able to persist in the medium and inside amoebas for 24 h, possibly because this is the optimal temperature and cultivation medium for promastigotes (de Oliveira Filho et al., 2024; Latifi and Salimi, 2020); although these conditions are not ideal for the cultivation of amoebas (Latifi and Salimi, 2020), the trophozoites can survive in the time necessary for the experimental observations.

Our results demonstrated visually the interaction of promastigotes and trophozoites and the prevalence of *L. amazonensis* inside amoeba was measured after 3 h of co-culture. At this timepoint,20.1 % of amoeba contained *L. amazonensis*, with an infection rate o 1.6 parasites per amoeba. The parasites were successfully isolated from trophozoites and differentiated in axenic promastigotes, demonstrating their viability inside amoeba.

The amoeba phagocytosed the promastigotes via the acanthopodia, which are spine-like structures on the surface used to capture particles(Siddiqui and Khan, 2012) as was revealed in the present study by the results obtained with optical, fluorescent, and electronic microscopy. These results corroborate the findings of Campo-Aasen and colleagues, who evaluated through transmission electron microscopy the interaction of *L. braziliensis* amastigotes isolated from ask in lesion with *A. castellanii* trophozoites the parasites were able to persist inside the trophozoites, and the amoeba extended acanthopodia within 3 h of co-cultivation (Campo-Aasen et al., 1988). Similar conclusions were obtained by Santos and colleagues, using optical microscopy (Santos et al., 2024).

Interestingly, most *L. amazonensis* promastigotes are able to interact with *A. castellanii* trophozoites through the flagellum, which also occurs when promastigotes infect cells of the mammalian mononuclear phagocyte system (Halliday et al., 2020; Martínez-López et al., 2018). Taken together, the results of previous studies, as well as the results of the present study, indicate that *Leishmania* can actively and passively penetrate the amoeba through phagocytosis in a manner comparable to the infection of mammalian macrophages (Ferreira et al., 2022; Martínez-López et al., 2018). The phagocytosed forms of *L. amazonensis* become rounded and lose their flagellum (amastigote-like forms) and resemble amastigotes the intracellular parasite form observed within the parasitophorous vacuoles of mammalian macrophages (Brioschi et al., 2022). Although it appears that these amastigotes-like forms are located within amoeba vacuoles, as previous pointed out by Santos and collaborators (Santos et al., 2024), only with the analysis of vacuole markers it is possible to confirm the localization of *Leishmania* inside amoebas.

Regarding the viability of both *Leishmania* and the amoeba after co-cultivation, the data indicated that in the first 3h both intracellular amastigote-likeforms and trophozoites were viable. No proliferation of *L. amazonensis* or amoeba was detected over longer periods. Indeed, over time, the number of amastigote-like forms decreased, and no parasites were found inside the *A. castellanii* trophozoites after 48 h. Indeed, this result corroborates the findings of Winiecka-Krusnell and collaborators, who also measured the percentage of amoebas with *L. tropica* for a period of 72 h of co-culture in PYG-W medium at 30°C; the portion of amoebas containing parasites was 78% and 6% at 3 and 24 h, respectively; and after 48 h no forms of *L. tropica* were found inside the trophozoites (Winiecka-Krusnell et al., 2009). *A. castellani* may act as a phagotrophic predator, have a well-developed phagosome-associated feeding pathway, and produce oxidative attack through ROS, and NO (Mungroo et al., 2021; Rayamajhee et al., 2021), which are microenvironments that can be harmful to intracellular *Leishmania* over time. However, preliminary data from our laboratory (data not shown) indicated no elevated levels of nitrite in the co-cultures of *A. castellanii* trophozoites and *L. amazonensis* promastigotes. Therefore, the lytic and killing mechanisms of *A. castellanii* during its interaction with *Leishmania* remain unknown.

Compared with mammalian macrophages, this kinetic profile, i.e. the infection rate decreased over time, was also observed in infected host cells isolated from murine lesions (Terreros et al., 2017), but not in the most commonly used cell systems murine peritoneal and bone marrow macrophage cultures, in which the number of intracellular amastigotes increased over time (96 h) (Santos-Pereira et al., 2019).

One of the objectives of this study was to evaluate whether *A. castellanii* could be used as a model organism for *in vitro s*tudies on *Leishmania*. These results demonstrate that *L. amazonensis* is indeed internalized by the amoeba; however, its survival rate decreased after 24 h. Thus, considering that the amoeba can clear *Leishmania* infection, this interaction could serve as an effective model of cellular leishmanicidal mechanisms that can be later compared to the macrophage-parasite relationship. Studies involving *A. castellanii* signaling pathways and cytolytic mechanisms during the period of interaction with *Leishmania*, combined with state-of-the-art techniques such as transcriptomics and proteomics, can reveal valuable information about constitutive molecules and/or molecules that are rapidly expressed with potent microbicidal functions, opening new possibilities for finding biologically active substances.

We do not know whether there is any ecological significance of the *A. castellani*-*Leishmania* interaction nor whether this occurs in its natural environment. However*, A. castellanii* acts as both a predator and environmental host for various pathogens and is considered the Trojan horse of the microbial world (Siddiqui and Khan, 2012). Thus, considering that the amoeba form of cystic resistance can be widespread in practically any environment, its trophozoite form may be associated with the presence of microbial biofilms (Lacerda and Lira, 2021; Mungroo et al., 2021; Zhang et al., 2023).*Acanthamoeba* spp. were isolated from wild mosquitoes *Aedes aegypti* (Otta et al., 2012);it is possible that the amoebas interact with the larval stage of the sandfly vector and, later, within the adult insect stage, interact with *Leishmania* promastigotes (Figure 6). The hypothesis put forward is that larval stages of sandflies ingest the amoebas found in aquatic environments and biofilms and acquire them in the digestive tract. When adult sandflies take a blood meal from mammals infected by *Leishmania*, they ingest infected cells, and the amastigotes transform into promastigote procyclics that interact with amoebas within the digestive tract (Figure 6). Although the viability of trophozoites in this microenvironment may be limited, a rapid interaction with *Leishmania* could exert some influence on *Leishmania*. Future studies are required to assess the role of *A. castellanii* as a natural host for different pathogens.

**Figure 6.**
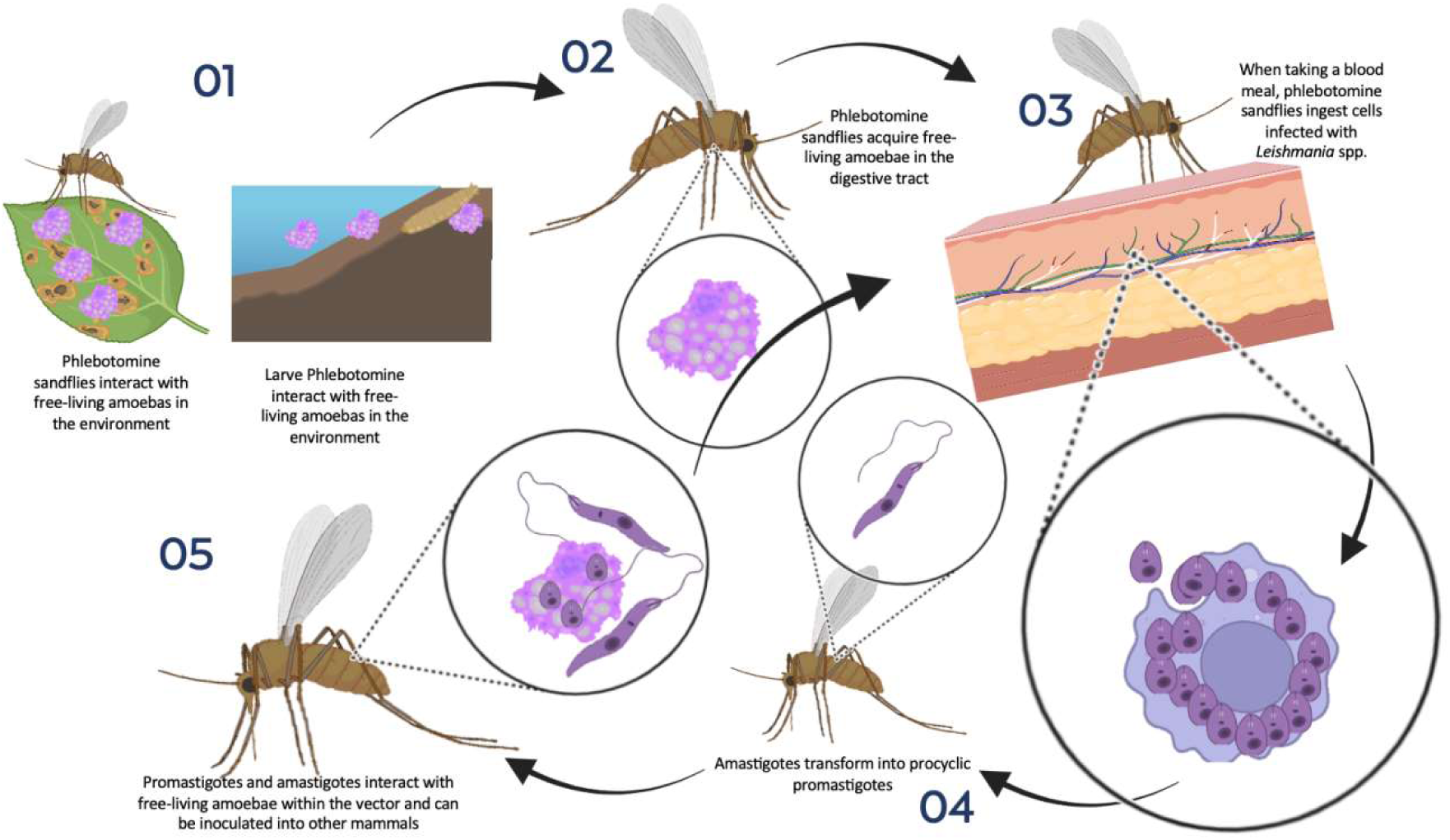
Hypothesis about the interaction between Free-Living Amoebas and *Leishmania* in the environment.

## 5. CONCLUSION

In this study, an understanding of the interaction between the amoeba *A. castellanii* and trypanosomatidae *L. amazonensis* was provided by combining quantitative kinetic analyses with optical, fluorescence, electronic, confocal, and live video microscopy. With the goal of establishing standard protocols for co-culture of promastigotes and trophozoites, we determined that the optimal experimental conditions, was RPMI medium + 10% SFB at 26 °C. *A. castellanii* internalizes *L. amazonensis* through its acanthopods, potentially via phagocytosis, and the promastigotes interact with trophozoites via their flagellum, which also occurs when parasites infect mammalian macrophages. *L. amazonensis* inside the amoeba become rounded and lose their flagellum (amastigote-like forms), similar to what occurs in amastigotes, the intracellular parasite form observed inside mammalian macrophages. Although these intracellular forms were successfully isolated from trophozoites after 3 h of co-culture and differentiated into axenic promastigotes, demonstrating their viability inside the amoeba, the percentage of amoebas with *L. amazonensis* was drastically reduced overtime and there were no amoebas with the parasites after 24 h of co-culture. Thus, considering that *A. castellanii* can clear *Leishmania* infection, this interaction could serve as an effective model of cellular leishmanicidal mechanisms that can be later compared with the macrophage-parasite relationship.

## Supplementary Material

**Figure 1.**
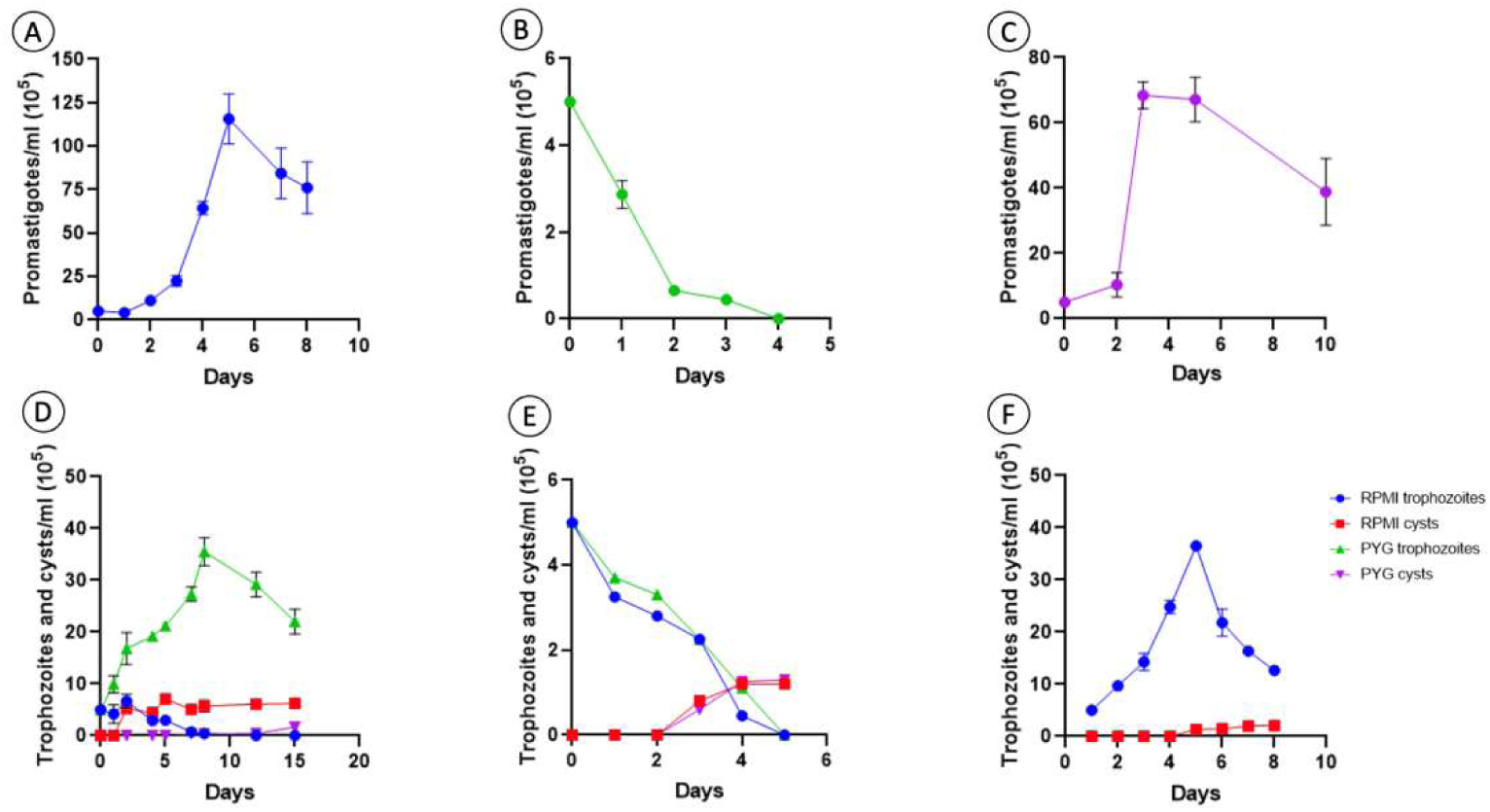
Proliferation curves of *L. amazonensis* promastigotes and *A. castellanii*. Approximately 5.10^5^ promastigotes/ml or trophozoites/ml were cultured and counted for 10 days. (A) RPMI + 10% SFB at 26°C; (B) PYG-W at 26°C (B) PYG-W+10% SFB at 26°C; (D) RPMI media + 10% SFB or PYG-W+ 10% SFB at 26°C. (E) RPMI media + 10% SFB or PYG-W+ 10% SFB at 37°C; (F) PYG-W + 10% of SFB at 34°C. Data are representative of three independent experiments.

**Figure 2.**
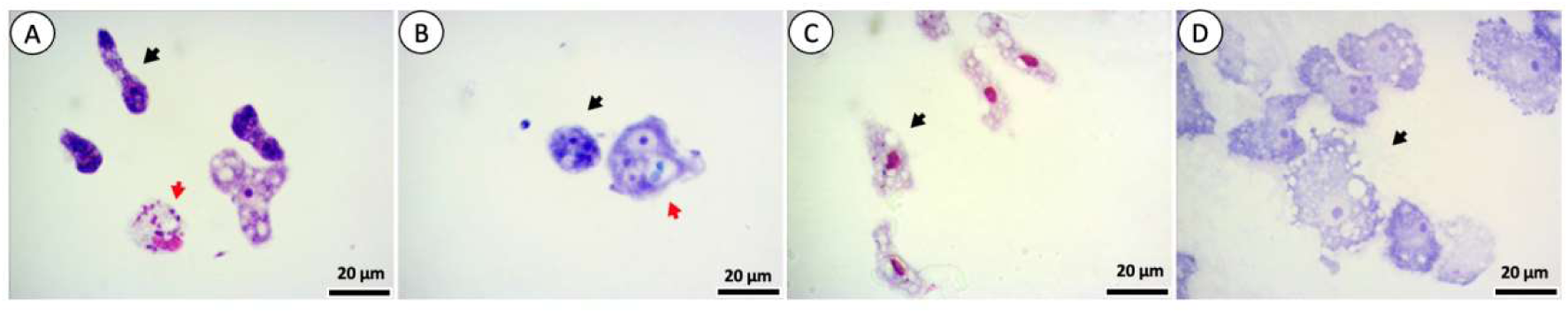
*A. castellanii* trophozoites in co-culture with *L. amazonensis* maintained for 3, 24, 48 and 72 hours. Red arrows indicate shapes intracellular cells of *L. amazonensis*, black arrows indicate trophozoites. Cells were stained with Giemsa. *A. castellanii* trophozoites (A-D).

**Figure 3.**
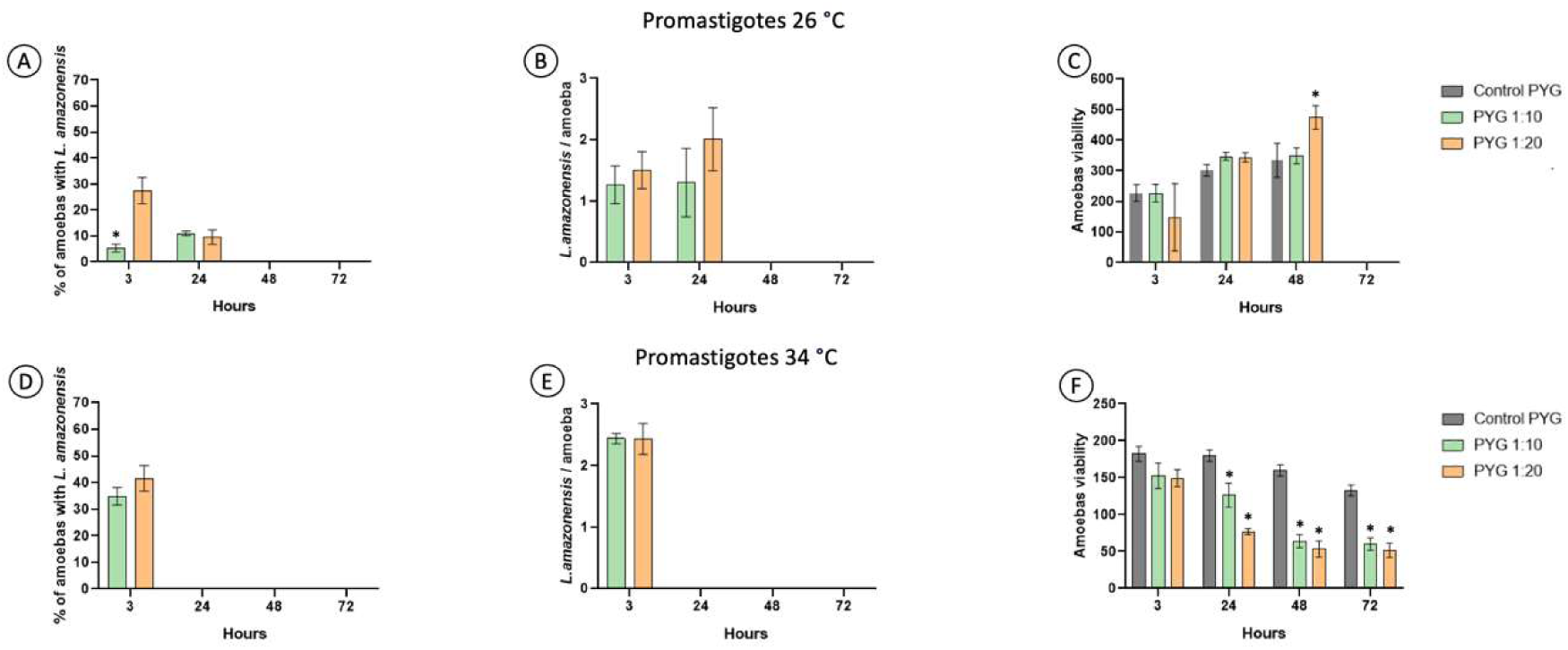
Interaction between *A. castellanii* trophozoites and *L. amazonensis.* Trophozoites and promastigotes were maintained at 26°C (A, B and C) and 34 °C (D, E and F) in PYG-W + 10% SFB media, 3, 24, 48 and 72 hours, with interaction ratios of 1:10 and 1:20 (trophozoite:promastigotes) and control (amoebas only). Data are representative of three independent experiments, performed in triplicate and values are expressed in mean ± SD; *p ≤ 0.05 (ANOVA).

**Figure 4.**
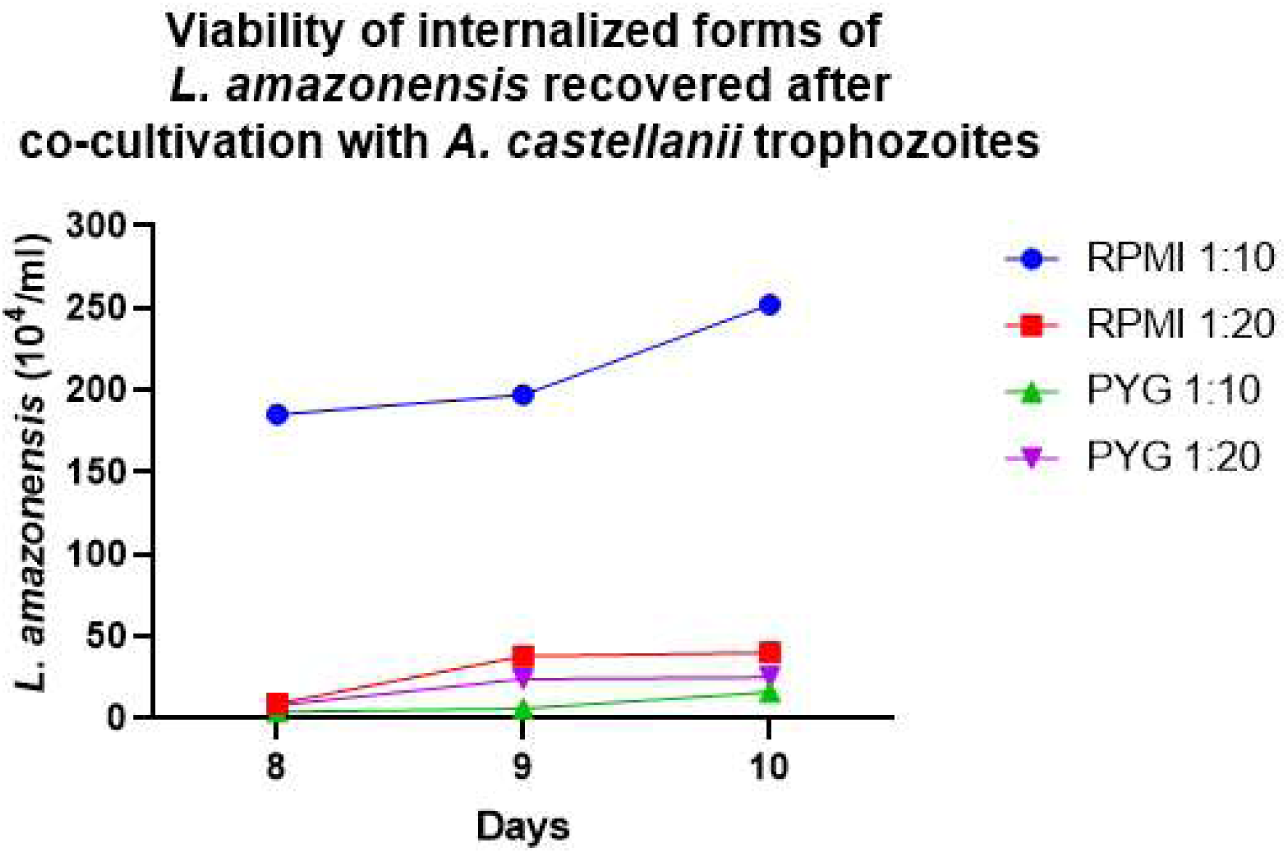
Proliferation curve of recovered intracellular forms of *L. amazonensis* after co-cultivation with *A. castellanii* in RPMI + 10% SFB or PYG-W + 10% SFB media, in interaction ratios 1:10 and 1:20 (trophozoite: promastigotes). Data are representative of three independent experiments.

**Figure 5.**
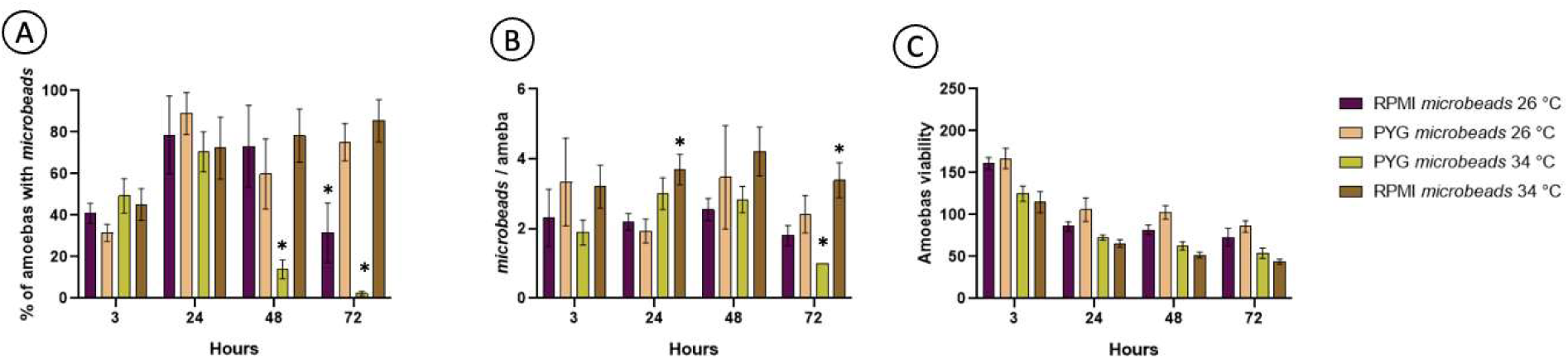
Interaction between *A. castellanii* trophozoites and microbeads. Trophozoites and promastigotes were maintained at 26°C and 34 °C (A, B, C, D, E and F) in RPMI + 10% SFB or PYG-W + 10% SFB medium, 3, 24, 48 and 72 hours. Data are representative of three independent experiments, performed in triplicate and values are expressed in mean ± SD; *p ≤ 0.05 (ANOVA).

**Figure 6.**
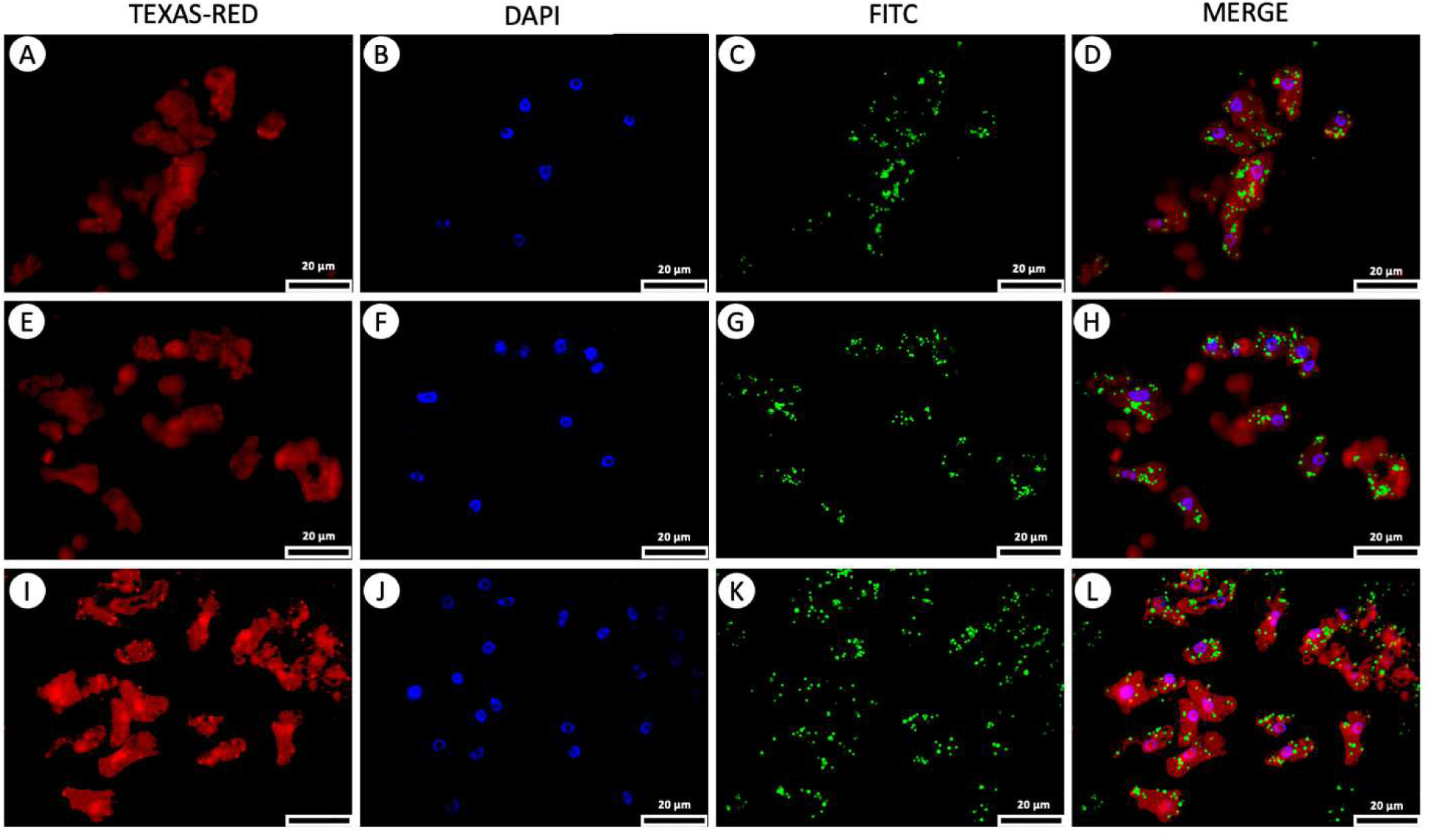
Fluorescence microscopy of trophozoites stained with DAPI (nucleus) and Celltracker (cytoplasm) interacting with microbeads. Celltracker(A, E and I). DAPI(B, F and J). FITC (*L. amazonensis*) (C, G and K). Merge (D, H and L).

**Figure 7.**
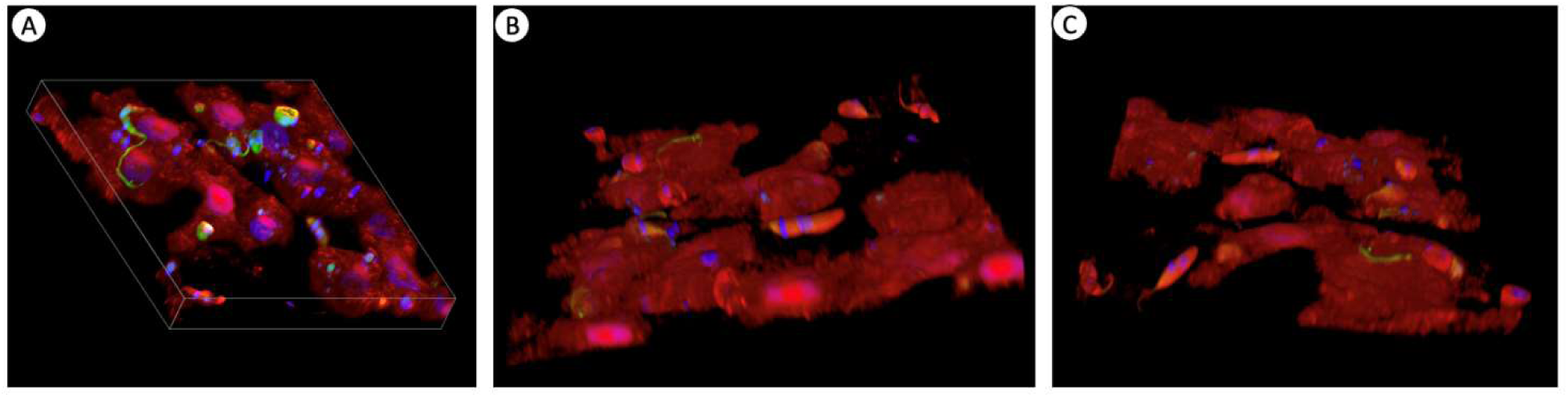
Three-dimensional volumetric (3D model) reconstruction of the sample obtained through confocal microscopyof trophozoites stained with DAPI and Celltracker interacting with *L. amazonensis* - GFP.

**Figure 8.**
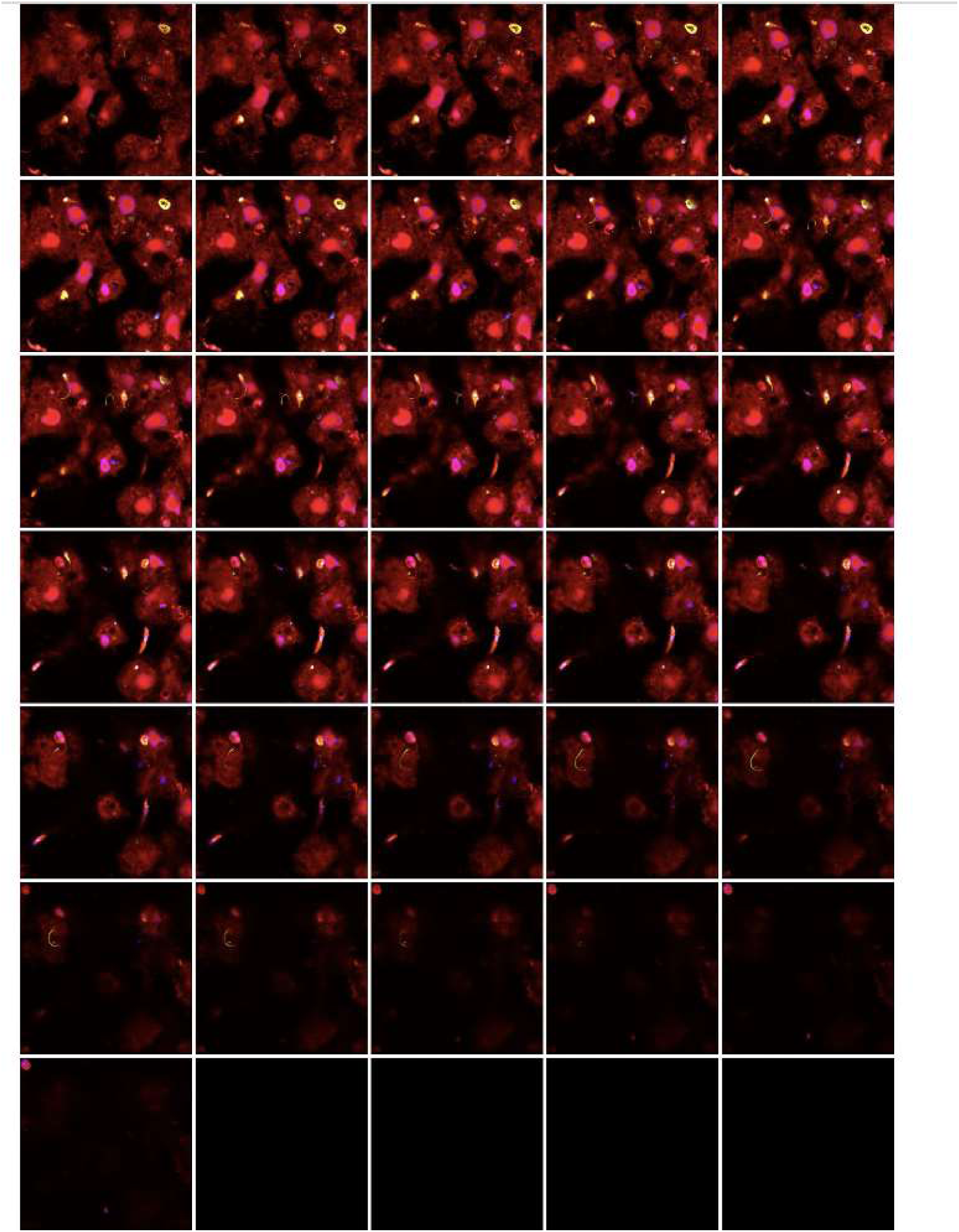
Plate of 31 views of parasite co-culture on Confocal microscopy.

**Figure 9.**
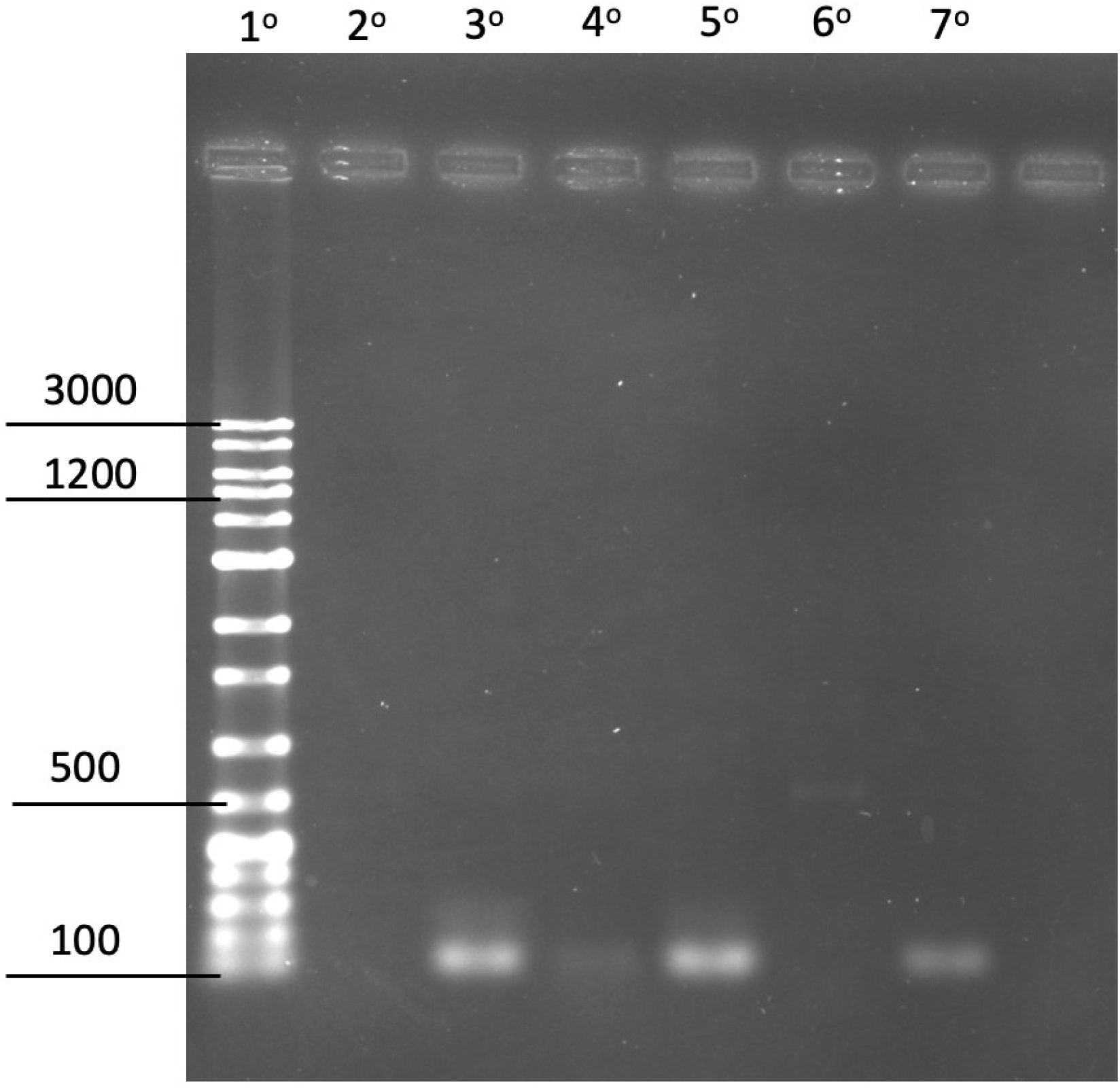
Validation of the oligonucleotide sequence (described in the methodology) for *L. amazonensis* and *A. castellanii*. 1^st^ line Molecular marker, 2^nd^ line *A. castellanii* trophozoites, 3^rd^ line *L. amazonensis* promastigotes, 4^th^ line Co-cultivation of *A. castellanii* trophozoites and *L. amazonensis* promastigotes (1:20), 5^th^ line *L. amazonensis* promastigotes, 6^th^ line *A. castellanii* trophozoites, 7^th^ line Co-cultivation of *A. castellanii* trophozoites and *L. amazonensis* promastigotes, 8^th^ line Negative control.

**Movie S1.** 3D visualization of the interaction between *A. castellanii* and *L. amazonensis* parasites.

**Movie S2.** Time-lapse video showcasing the dynamic interaction between *A. castellanii* and *L. amazonensis* parasites over a 24h period. Observe the movement, behavior, and possible phagocytosis events in this captivating micro-scale interaction.

**Movie S3.** Time-lapse video showcasing the dynamic interaction between *A. castellanii* and *L. amazonensis* parasites for a brief moment.

## Supporting information

### Funding

This study was funded by Conselho Nacional de Desenvolvimento Científico e Tecnológico (CNPq, 304309/2021-4) and Fundação de Amparo a Pesquisa do Estado de São Paulo (FAPESP, 2018/23302-6). Leonardo F. Geres was recipient of Coordenação de Aperfeiçoamento de Pessoal de Nível Superior Fellowships (CAPES, 88887.713017/2022-00). Selma Giorgio and Marcelo Brocchi are CNPq research productivity fellows.

### Credit authorship contribution statement

**Leonardo F. Geres:** Methodology, Investigation, Data curation, Writing – original draft. **Pedro Henrique Gallo Francisco:** Methodology, Investigation. **Diullia de Andrade Machado:**Methodology, Investigation. **Francisco Breno Silva Teófilo:** Methodology, Investigation. **Marcelo Brocchi:** Investigation. **Selma Giorgio:** Conceptualization, Validation, Visualization, Supervision, Writing – review & editing.

### Declarations

All authors disclose that they do not have any financial and personal relationships with other people or organizations that improperly influence(bias) our work.

## Acknowledgments

We would like to thank the access to equipment and assistance provided by the Electron Microscope Laboratory (LME/UNICAMP).We thank the access to equipment and assistance provided by the National Institute of Science and Technology on Photonics Applied to Cell Biology (INFABIC) at the State University of Campinas; INFABIC is co-funded by Fundação de Amparo a Pesquisa do Estado de São Paulo (FAPESP) (2014/50938-8) and Conselho Nacional de Desenvolvimento Cientifico e Tecnológico (CNPq) (465699/2014-6).We acknowledge Editage (www.editage.com) for their support in improving the English language quality of this manuscript. We thank Victor Agostino for his careful review of the manuscript.

## Notes

### Competing Interest Statement

The authors have declared no competing interest.

